# Estimating energy expenditure from wrist and thigh accelerometry in free-living adults: a doubly labelled water study

**DOI:** 10.1101/370247

**Authors:** Tom White, Kate Westgate, Stefanie Hollidge, Michelle Venables, Patrick Olivier, Nick Wareham, Soren Brage

## Abstract

**Background:** Many large studies have implemented wrist or thigh accelerometry to capture physical activity, but the accuracy of these measurements to infer Activity Energy Expenditure (AEE) and consequently Total Energy Expenditure (TEE) has not been demonstrated. The purpose of this study was to assess the validity of acceleration intensity at wrist and thigh sites as estimates of AEE and TEE under free-living conditions using a gold-standard criterion.

**Methods:** Measurements for 193 UK adults (105 men, 88 women, aged 40-66 years, BMI 20.4-36.6 kg·m^-2^) were collected with triaxial accelerometers worn on the dominant wrist, non-dominant wrist and thigh in free-living conditions for 9-14 days. In a subsample (50 men, 50 women) TEE was simultaneously assessed with doubly labelled water (DLW). AEE was estimated from non-dominant wrist using an established estimation model, and novel models were derived for dominant wrist and thigh in the non-DLW subsample. Agreement with both AEE and TEE from DLW was evaluated by mean bias, Root Mean Squared Error (RMSE) and Pearson correlation.

**Results:** Mean TEE and AEE derived from DLW was 11.6 (2.3) MJ·day^-1^ and 49.8 (16.3) kJ·day^-1^·kg^-1^. Dominant and non-dominant wrist acceleration were highly correlated in free-living (r=0.93), but less so with thigh (r=0.73 and 0.66, respectively). Estimates of AEE were 48.6 (11.8) kJ·day^-1^·kg^-1^ from dominant wrist, 48.6 (12.3) from non-dominant wrist, and 46.0 (10.1) from thigh; these agreed strongly with AEE (RMSE ~12.2 kJ·day^-1^·kg^-1^, r ~0.71) with small mean biases at the population level (~6%). Only the thigh estimate bias was statistically significantly different from the criterion. When combining these AEE estimates with estimated REE, agreement was stronger with the criterion (RMSE ~1.0 MJ·day^-1^, r ~0.90). Conclusions: In UK adults, acceleration measured at either wrist or thigh can be used to estimate population levels of AEE and TEE in free-living conditions with high precision.

## Introduction

Characterising the energy balance of individuals in free-living conditions requires an accurate assessment of total energy expenditure. Total energy expenditure can be measured with high precision using the doubly labelled water technique^1^ but this is an expensive undertaking that requires elaborate sample collection and analysis infrastructure, making it less feasible for large-scale deployment or application in clinical settings. In most people, the largest component of total energy expenditure is resting energy expenditure, which can be predicted from anthropometric information with reasonable accuracy^2,3^. Diet-induced thermogenesis is less variable and ordinarily constitutes approximately 10% of total energy expenditure^4^. The predominant source of uncertainty in total energy expenditure estimates is the highly-variable activity energy expenditure component, which has proven difficult to capture by subjective instruments such as questionnaires^5,6^. Body-worn sensors such as accelerometers have the potential to provide a relatively cheap and reliable solution to this problem^7^, if valid inference models can be devised to estimate activity energy expenditure from the measurements they record.

In recent years, wrist-worn accelerometers have become a popular measurement modality for objectively capturing free-living physical activity in large-scale studies^8–10^. Devices worn on the wrist are generally considered to be less burdensome for participants than those worn on other anatomical sites^11^. This has led to improved wear protocol adherence and thus to measurements with potentially greater representation of habitual physical activity levels. However, despite their recent increase in popularity, their utility in the estimation of activity energy expenditure has yet to be tested against gold-standard techniques in a sufficiently large sample of men and women in free-living^12^. Furthermore, some large studies ^8–10^ have committed to measuring only one of either the dominant wrist or non-dominant wrist, and the relationship between these two measurements also remains understudied.

In previous work, we derived parametric models to estimate activity energy expenditure intensity from non-dominant wrist acceleration (reproduced in Table 2) using a dataset (n=1050) of simultaneous non-dominant wrist and individually-calibrated combined heart rate and movement sensing signals collected under free-living conditions^13^. We evaluated the models in a large holdout sample (n=645) and found that they explained 44-47% of the variance in activity energy expenditure with no significant mean bias at the population level. However, as this comparison was against a silver-standard measurement of activity volume, these estimation models could be more conclusively validated by integrating the estimated activity energy expenditure signal over time, and assessing agreement of activity volume with a gold-standard criterion such as doubly labelled water. This approach has been used to validate combined heart rate and movement sensing ^14–16^ against which the models were originally derived.

Thigh-worn devices have typically been employed in smaller studies to measure time spent in a sitting posture, in order to infer sedentary time. This is possible because the distribution of gravity over the three axes can be interpreted using a simple equation to calculate thigh inclination. However, thigh acceleration has received comparatively little attention as a measure of physical activity intensity, though it features prominently in activity classification experiments^17^. In epidemiological settings, thigh-worn sensors have been complemented by other sensors with the intention to capture physical activity separately^18^.

The primary aim of this study was to describe the absolute validity of a previously established activity energy expenditure prediction model ^13^ when applied to both wrists, and to evaluate the validity of this estimation in predicting total energy expenditure when combined with a simple anthropometric prediction of resting energy expenditure^2^. The second aim was to use the same approach to derive and validate similar energy expenditure estimation models using thigh acceleration. The third aim was to explore the relationship between the dominant wrist, non-dominant wrist and thigh acceleration measures in free-living, and to derive intensity models to facilitate harmonisation.

## Subjects and Methods

Participants were recruited from the Fenland study, an ongoing cohort described in detail elsewhere ^19^. We aimed to recruit participants who had previously indicated that they were interested in participating in future studies, were aged between 40 and 70 years, with a BMI between 20 and 50 kg·m^-2^. Recruitment aimed to balance age, sex and BMI distributions. Participants were invited to attend an assessment centre on two separate occasions, separated by a free-living period of 9 to 14 days. Ethical approval for the study was obtained from Cambridge University Human Biology Research Ethics Committee (Ref: HBREC/2015.16). All participants provided written informed consent.

Weight was measured to the nearest 0.1 kg using calibrated digital scales (TANITA model BC-418 MA; Tanita, Tokyo, Japan) at both visits. Height was measured to the nearest 0.1 cm using a stadiometer (SECA 240; Seca, Birmingham, UK) at the first clinic visit. Body composition was also measured by DXA (Lunar Prodigy Advanced, GE Healthcare, USA) as part of the Fenland study.

Total energy expenditure was measured by doubly labelled water in 100 of the participants. Prior to the first clinic visit, participants self-reported their current weight, which was used to provide a body-weight specific dose of ^2^H_2_^18^O (70 mg ^2^H_2_O and 174 mg H_2_^18^O per kg body weight). Participants brought a baseline urine sample to their first clinic visit, and a second baseline sample was taken at the clinic visit, prior to dosing. Participants were provided labelled sampling bottles and asked to collect one urine sample per day for the next 9-10 days, at a similar time each day but not the first void of the day. Participants were asked to record the date and time of each measurement on the sample bottle label and separately on a provided timesheet. Participants were asked to store the samples in a container in a cool, dry place, such as a refrigerator, and to return those samples at their second clinic visit at the end of their free-living measurement period. Isotope ratio mass spectrometry (^2^H, Isoprime, GV Instruments, Wythenshaw, Manchester, UK and ^18^O, AP2003, Analytical Precision Ltd, Northwich, Cheshire, UK) was used to measure the isotopic enrichment of the samples. All samples were measured alongside laboratory reference standards, previously calibrated against the international standards Vienna-Standard Mean Ocean Water (vSMOW) and Vienna-Standard Light Antarctic Precipitate (vSLAP) (International Atomic Energy Agency, Vienna, Austria). Sample enrichments were corrected for interference according to Craig ^20^ and expressed relative to vSMOW. Rate constants and pool sizes were calculated from the slopes and intercepts of the log-transformed data, with total CO_2_ production (RCO_2_) calculated using the multi-point method of Schoeller ^21^. RCO_2_ was converted to total energy expenditure ^22^ where the respiratory quotient was informed by the macronutrient composition of the diet (see below).

Resting metabolic rate was measured at the start of both clinic visits during a fifteen-minute rest test by respired gas analysis (OxyconPro, Jaeger, Germany). A seven-breath running median was calculated and the lowest observed average rate over a five minute consecutive window was found, which was scaled down by 6% to compensate for within-day elevation of resting metabolic rates ^23^. Basal metabolic rate was also estimated via three different equations which differ in the specific body composition information utilised ^2,24,25^. Resting energy expenditure was primarily characterised as the nearest measured value to the mean average estimated value, and a further sensitivity analysis was conducted using exclusively measured values. The final 24-hour resting energy expenditure estimates also included an adjustment for a 5% lower metabolic rate during sleep^26^, according to their reported mean sleep duration.

At the second clinic visit, participants were asked to complete a Food Frequency Questionnaire^27^, which was used to estimate dietary intake over the past year. The food frequency data was processed using FETA^28^, and the resulting calorie-weighted macronutrient profile was used to calculate the Food Quotient and diet-induced thermogenesis^29^. Diet-induced thermogenesis was normalised by the total energy expenditure to total energy intake ratio, as done previously^14^.

At the first clinic visit, participants were fitted with three waterproof triaxial accelerometers (AX3, Axivity, Newcastle upon Tyne, UK); one device was attached to each wrist with a standard wristband, and one was attached to the anterior midline of the right thigh using a medical-grade adhesive dressing. The devices were setup to record raw, triaxial acceleration at 100 Hz with a dynamic range of ±8 g (where g refers to the local gravitational force, roughly equal to 9.81 m·s^-2^). Participants were asked to wear them continuously for the following 8 days and nights whilst continuing with their usual activities. They were also asked to record their main sleep using a sleep diary throughout the free-living period.

The signals were resampled from their original irregularly timestamped intervals to a uniform 100 Hertz signal by linear interpolation, and then calibrated to local gravity using a well-established technique^30,31^, without adjustment for temperature changes within the record. Periods of nonwear were identified as windows of an hour or more wherein the device was inferred to be completely stationary ^11^, where stationary is defined as standard deviation in each axis not exceeding the approximate baseline noise of the device itself (10 milli-g). Vector Magnitude (VM) was then calculated from the three axes (VM (X,Y,Z) = (X^2^ + Y^2^ + Z^2^)^0.5^), from which two acceleration intensity metrics were derived ^32^; Euclidean Norm Minus One (ENMO) subtracts 1 g from VM and truncates any negative results to 0, and High-Pass Filtered Vector Magnitude (HPFVM) applies a fourth-order high-pass filter to the signal at a 0.2 Hertz cut-off (3 dB). These analyses were performed using pampro v0.4.0^33^.

In the non-doubly labelled water group (n=93), multi-level linear regression with random effects at the participant level was used to characterise each of the pairwise relationships between dominant wrist, non-dominant wrist and thigh acceleration intensity using synchronised 5-minute level data from each source. We used these intensity relationships to derive new activity energy expenditure estimation models for thigh and dominant wrist-worn devices, by substituting the non-dominant wrist term in our original models with the derived equation to harmonise either dominant wrist or thigh acceleration to non-dominant wrist acceleration.

Activity energy expenditure was estimated separately from each of the acceleration signals by directly applying the appropriate linear and quadratic equations given in Table 2 to 5-second level data; the resulting 5-second level estimated activity energy expenditure signal was then summarised to a mean-per-day average activity energy expenditure using diurnal adjustment to compensate for any between-individual bias introduced by periods of nonwear^34^. To ensure a stable estimate of this circadian model, a minimum of 72 hours of valid data was required per signal to be included in the analyses. Predicted total energy expenditure (in MJ·day^-1^) was calculated as the sum of predicted activity energy expenditure and predicted resting energy expenditure from the simplest model (using only age, sex, height and weight)^2^, and dividing the result by 0.9 to account for diet-induced thermogenesis^4^. Agreement between these two predictions against measured activity energy expenditure and total energy expenditure from doubly labelled water was formally tested by calculating the pairwise mean bias and 95% limits of agreement, Root Mean Squared Error (RMSE) and Pearson’s correlation coefficient. Linear regression was used to characterise the relationship between the acceleration measurements and activity energy expenditure/total energy expenditure derived from doubly labelled water. As the main focus of this paper is on absolute validity, these relative validity results are supplied in the supplementary material.

The statistical tests were performed using Python v3.6 and Stata v14 (StataCorp, TX, USA).

## Results

A descriptive summary of participant characteristics is given in Table 1. We recruited 193 participants, and the group measured by doubly labelled water was split equally between men and women. According to the doubly labelled water measurements, mean (standard deviation) total energy expenditure was 11.6 (2.3) MJ·day^-1^, of which 6.6 (1.2) MJ·day^-1^ was resting energy expenditure. Mean (standard deviation) activity-related acceleration (ENMO) per day was 32.4 (8.3) milli-g on the dominant wrist, 28.8 (7.7) milli-g on the non-dominant wrist, and 27.8 (10.9) milli-g on the thigh. Mean dominant wrist acceleration was higher than non-dominant wrist in 84% of participants.

Some accelerometry measurements were not included in the analyses due to a combination of devices being lost by participants (n=7), device failures (n=3), user error upon download (n=3), and insufficient wear time (n=3). Of those files that overlapped with doubly labelled water measurements, 3 were dominant wrist records, 3 were non-dominant wrist and 9 were thigh records. There was no loss of data in the doubly labelled water, anthropometry or food frequency questionnaire measurements.

Table 2 lists the derived equations to predict activity energy expenditure from each of the sensors, as informed by the harmonisation equations which are supplied in Supplementary Table 1. For brevity, Table 3 summarises the absolute validity of the quadratic HPFVM models applied to measurements from both wrists and thigh with respect to activity energy expenditure, and Table 3 summarises agreement with total energy expenditure derived from doubly labelled water. A Bland-Altman plot illustrating the agreement of these estimates is supplied in Figure 1. A table summarising the remaining models is given in Supplementary Table 2.

The difference in performance between each estimation model was very minor; all activity energy expenditure estimates had small negative mean biases (underestimates) at the population level (average −2.8 kJ·day^−1^·kg^−1^) but of these only the thigh model biases were statistically significant. RMSEs for activity energy expenditure ranged from 11.9 to 13.5 kJ ·day^−1^·kg^−1^ (24 to 27% of the mean), and 1.0 to 1.2 MJ·day^−1^ for total energy expenditure (8 to 10% of the mean). Pearson correlations ranged from 0.6 to 0.69 with activity energy expenditure, and from 0.87 to 0.91 with total energy expenditure. Combined estimates using two or more sensors lead to very negligible performance improvements over single-sensor estimates. Signed estimation errors were nominally positively correlated with body fat percentage when using our primary characterisation of resting energy expenditure (r=0.18- 0.25), and less so with exclusively measured values (r=0.10-0.17).

In the non-doubly labelled water group, 88 participants had at least 3 days of valid simultaneous wrist signals during free-living, and 84 had simultaneous wrist and thigh signals; around 200 000 5-minute observations included in each of the regression analyses. The between-individual explained variance between dominant and non-dominant wrist intensity signals was approximately 86% (99% within-individual), and the average between-individual explained variance between wrist and thigh intensities was approximately 49% (97% within-individual). The derived linear models to harmonise the acceleration signals are listed in Supplementary Table 1. The final models given to estimate activity energy expenditure from dominant wrist and thigh in Table 2 were the result of substituting these harmonisation equations into the original non-dominant wrist models.

## Discussion

In this work, we have applied our previously derived activity intensity estimation models ^13^ to wrist acceleration signals (after harmonising the intensity of dominant wrist to non-dominant wrist) and investigated their agreement with a gold-standard measure of activity energy expenditure. We arrived at estimates that were highly correlated with the criterion (r > 0.6) with small and non-significant mean biases at the population level from both wrists and low RMSEs of approximately 12 kJ·day^-1^·kg^-1^. We have also introduced and validated new intensity estimation models for thigh acceleration, demonstrating similar performance to the wrist models. We observed that dominant wrist acceleration was on average 12% higher than non-dominant wrist in free-living individuals, but that those measures were very highly correlated (r=0.93), allowing us to derive conversion models which harmonise acceleration intensity measured at either wrist. To our knowledge, this is the first demonstration of the absolute validity of a time-integrated predictive model of activity intensity for either wrist or thigh accelerometry.

Our findings on the high correlation between dominant wrist and non-dominant wrist acceleration in free-living individuals are consistent with a previous study in a small convenience sample (n=40)^35^. They also observed ~5% higher dominant wrist than non-dominant wrist acceleration, but it was not a statistically significant difference, perhaps due to the shorter duration of measurement and smaller sample size. In our relative validity tests, we found that each wrist separately explained a similar variance in activity energy expenditure, and inclusion of both wrist measurements in the linear models did not drastically improve performance over either wrist measurement alone. Taken together, these results are indicative of a high degree of upper-body symmetry. One implication of these findings is that irrespective of hand dominance, wrist acceleration measurements are naturally conducive to harmonisation across studies, making them well suited to pooled- and meta-analysis.

Conversely, it implies that implementing dual wrist measurements may be a largely redundant exercise for studies whose primary intention is to capture activity energy expenditure. However, there is a possibility that future methodological advances in the field of activity recognition may be able to better utilise simultaneous wrist signals, which could yield a more precise instantaneous estimation of activity energy expenditure.

The estimation models validated herein for the wrist were derived using a training dataset in which non-dominant wrist acceleration data was collected at 60 Hz with a GeneActiv device ^13^, and were successfully validated using 100 Hz data collected with an Axivity AX3. With an additional harmonisation step, the model also translated to acceptably strong inferences on the dominant wrist, albeit with a slightly increased error. This indicates that our models capture a generalized biomechanical relationship of wrist movement, rather than being superficial transformations of a specific device’s output to activity energy expenditure. It therefore suggests that these models are applicable to any wrist-worn device which provides raw, unfiltered triaxial acceleration data expressed in SI units.

The associations between wrist acceleration and observations from DLW have been reported before, in pregnant and non-pregnant Swedish women ^11^. In that population it explained 27% of the variance in activity energy expenditure (kJ·day^-1^·kg^-1^) in non-pregnant women (n=48), but only 5% in pregnant women (n=26); however, those wrist measurements were evenly divided between left and right wrist, which most likely lead to a mix of dominant and non-dominant wrist measurements and potentially attenuated the correlations.

The previously established estimation models applied to the non-dominant wrist resulted in robust estimates with small, non-significant mean biases, which is a strong justification for using this inference scheme to infer activity energy expenditure in free-living individuals. The higher average of the dominant wrist would have led to a significant overestimation had we applied the original non-dominant wrist model, but our harmonisation approach effectively scaled the dominant wrist measure down to the level of non-dominant wrist, ultimately leading to virtually identical results. We note that physical activity was measured by dominant wrist accelerometry in UK Biobank^8^. We have now demonstrated the validity of this approach in a demographically comparable sample. Specifically, the absolute validity result for ENMO in Supplementary Table 2 demonstrates that our linear estimation model applied to ENMO at 5-second resolution yielded a valid activity energy expenditure estimate, with a small mean bias and a RMSE of 13 kJ·day^-1^·kg^-1^ and high correlation (r=0.61).

Consequently, we can use the equations for dominant wrist in Table 2 to solve for salient energy expenditure values – for example, 3 metabolic equivalents (activity energy expenditure ~142 J·min^-1^·kg^-1^) is the generally accepted threshold for “moderate” activity intensity, and our ENMO equations suggest this is approximately 159 milli-g on the dominant wrist.

Our findings for the thigh acceleration models demonstrate that thigh-worn accelerometers capture an information-rich biomechanical signal, from which valid estimates of activity energy expenditure can be made. As a consequence of the larger y-intercepts of the thigh models, their minimum estimated activity energy expenditure ranges from 10 to 18 J·min^-1^·kg^-1^ (0.15-0.25 metabolic equivalents). To our knowledge, only one previous study has described the association between thigh acceleration and activity energy expenditure from doubly labelled water, in a small study of free-living cancer patients and controls^36^; which reported very low agreement between the manufacturer’s proprietary activity energy expenditure prediction and the criterion. While thigh-worn sensors do not yet have the same popularity as wrist-worn sensors^37,38^, large-scale data collections are planned for the future^39^. Our models enable new analyses to be conducted in those existing datasets, and may make thigh-worn accelerometry a more appealing option for future studies if issues of feasibility can be addressed.

Some have suggested that simple movement intensity approaches should be replaced by more sophisticated models that utilise a broader range of signal features^40,41^. Recent efforts to estimate energy expenditure have utilised a range of machine learning approaches, such as neural networks ^42–44^ and random forests^40^. While we are not aware of any such methodology with a performance that exceeds the simpler models validated in this paper, this is an interesting area of future work.

The results of our absolute validity tests demonstrate that deriving intensity models using a “silver-standard” criterion (such as individually-calibrated heart rate and uniaxial movement sensing) in a large sample of free-living adults is a sound approach. The combined sensing estimate of activity energy expenditure is less precise than respiratory gas analysis which can be captured in laboratory studies ^45^ but there are several reasons why we have been able to derive superior models to previous approaches. Firstly, the dataset was collected in free-living participants, and is therefore representative of the intended application, as opposed to artificial scenarios and activities performed in a laboratory. Secondly, the combined sensing approach embedded in a cohort study allowed the collection of a volume of data many orders of magnitude greater than any laboratory study has for this purpose. Our training dataset alone contained over 16.6 person-years of observation (>1.7 million data points). One disadvantage of this approach is that we are unable to capture categorical labelled data, so there is no opportunity to explore activity type recognition.

It is appropriate to compare our absolute validity results here with those of combined sensing itself ^14^. The best estimate with treadmill test calibration resulted in a RMSE of 20 kJ·day^-1^·kg^-1^ (30% of the 66 kJ·day^-1^·kg^-1^ criterion mean), non-significant positive mean bias of approximately 4 kJ·day^-1^·kg^-1^ (6%) at the population level, and a correlation of 0.67 in a sample of 50 UK adults. Compared to the present results, all estimations here had considerably lower RMSEs of around 12 kJ·day^-1^·kg^-1^ (25% of the 50 kJ·day^-1^·kg^-1^ mean), similar magnitude but negative mean biases (~6%), but generally higher correlations. However, our study participants were significantly less active overall according to the criterion, ultimately leading to a similar relative accuracy. Combined sensing model errors were also uncorrelated to body fat percentage, whereas errors of accelerometry-only models seem to display this characteristic, albeit less so in the present study (r=0.22 versus r=0.63 for uniaxial trunk acceleration). Contrasting the feasibility of the methods, however, wrist accelerometry has the advantages of being cheaper, less burdensome to both participants and research staff, and does not require individual calibration using an exercise test. Comparing performance of other devices worn on the upper limbs, validation of the now-discontinued SenseWear Pro3 and Mini also achieved no significant bias with respect to total energy expenditure, but with lower correlations (r=0.84) than any of our total energy expenditure models (r=0.9) and wider limits of agreement ^46^ and with lower feasibility.

In summary, we have evaluated the absolute validity of intensity models of activity energy expenditure from wrist and thigh accelerometry, and concluded that they provide precise and accurate estimates in free-living adults. With the addition of predicted resting energy expenditure to produce total energy expenditure, we found even stronger validity at the population level. Considering its feasibility, wrist accelerometry emerges as a viable candidate for deployment in a large scale studies, including physical activity surveillance and the prediction of total energy expenditure in dietary surveys.

## Acknowledgements

We are very grateful to the participants who took part in this study. We thank the principal investigators of the Fenland study for allowing us to recruit from this study population, and the functional teams of the MRC Epidemiology Unit (Study Coordination, Field Epidemiology, Anthropometry, Data Management, IT) for supporting the study. We would like to specifically acknowledge Lewis Griffiths, Katie Palmer, and Eoin McNamara for their assistance in the data collection for this study, and Annie Schiff, Richard Salisbury and Nicola Kimber for study co-ordination and recruitment.

We would like to thank Eirini Trichia from the MRC Epidemiology Unit for processing the FFQ data with the FETA package. We would also like to thank the stable isotope team from the MRC Elsie Widdowson Laboratory: Priya Singh, Elise Orford and Kevin Donkers for the DLW preparation and analysis.

Medical Research Council and UK Biobank are acknowledged for covering costs of the fieldwork. Newcastle University and MedImmune are acknowledged for covering the costs of the doubly labelled water measurements.

## Abbreviations list

Activity Energy Expenditure (AEE)
ADoubly-Labelled Water (DLW)
ADiet-Induced Thermogenesis (DIT)
AEuclidean Norm Minus One (ENMO)
AFood Quotient (FQ)
AFood Frequency Questionnaire (FFQ)
AHigh-pass Filtered Vector Magnitude (HPFVM)
AResting Energy Expenditure (REE)
Total Energy Expenditure (TEE)
Vector Magnitude (VM)

---

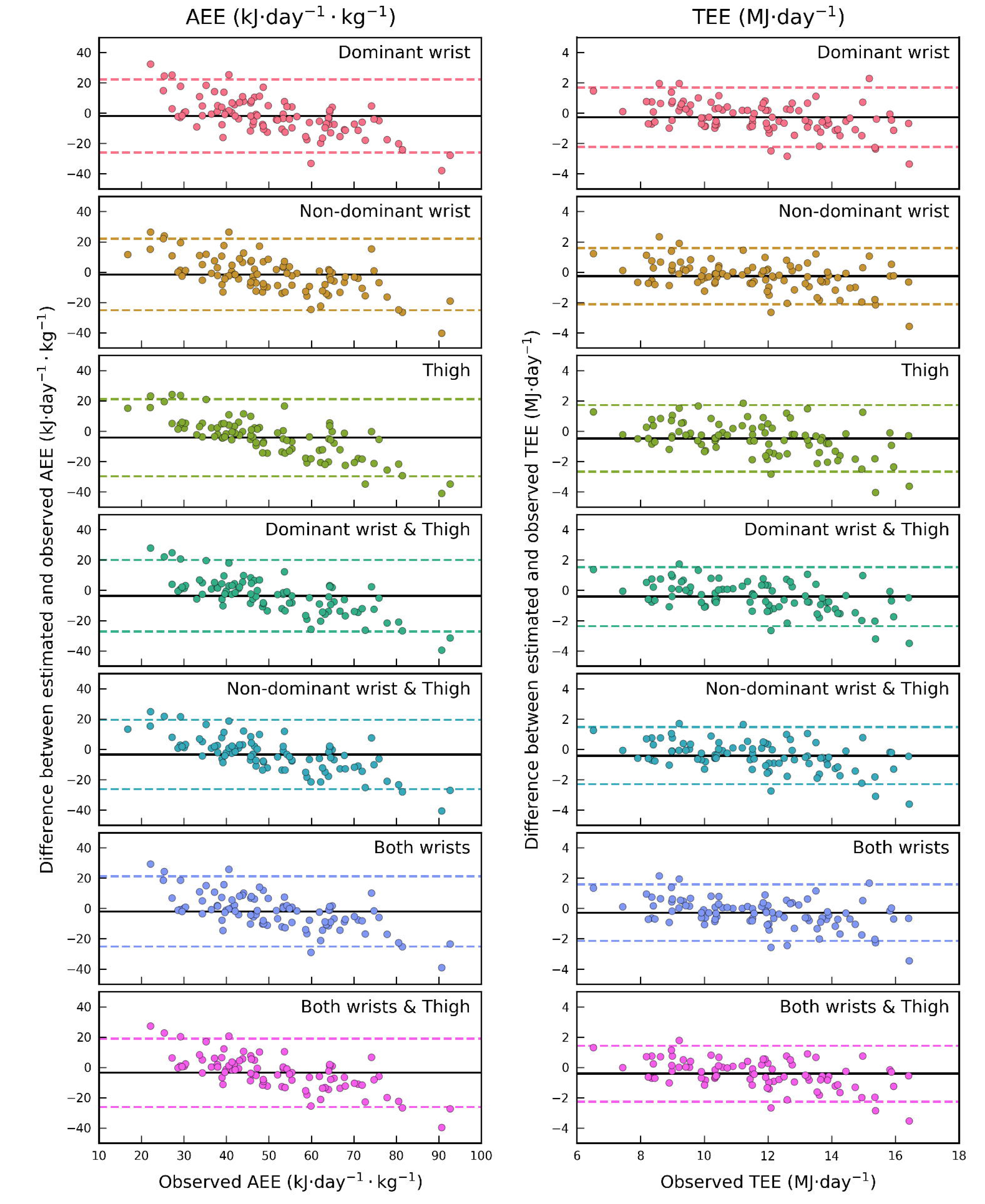
Participant characteristics, provided separately for the doubly labelled water and non-doubly labelled water groups. Derived linear and quadratic equations to estimate activity energy expenditure (J·min^-1·^kg^-1^) from wrist and thigh acceleration intensity. (4.184 J · min^-1·^kg^-1^ = 1 cal, and 71.225 J·min^-1·^kg^-1^ = 1 net Metabolic Equivalent Task (MET)). Agreement between estimated activity energy expenditure from the HPFVM quadratic models with those derived from doubly labelled water. An asterisk (*) next to a bias value indicates statistical significance according to a paired t-test (p < 0.05). Bland-Altman plots illustrating agreement between the activity energy expenditure and total energy expenditure estimates from HPFVM Quadratic models with those from doubly labelled water, where the X-axis indicates the observed values.

